# SWI3 regulates male sex determination in *Marchantia polymorpha*

**DOI:** 10.1101/2025.10.29.685095

**Authors:** Maria Lozano-Quiles, Parth K. Raval, Stefan A Rensing, Sven B. Gould

## Abstract

Land plants alternate between multicellular haploid and diploid phases, a life cycle requiring tight coordination between vegetative growth and sexual reproduction. Chromatin regulators such as SWI3, a core subunit of the SWI/SNF complex, are ancestral and highly diversified in angiosperms. As their functions outside of angiosperms remain unclear, we investigated *SWI3* of the liverwort *Marchantia polymorpha*, a non-vascular dioecious land plant. A mutation in the promoter of Mp*SWI3* affected gametangiophore development and spermiogenesis in males, revealing its male-specific role in reproductive development. The Mp*SWI3* mutant line amplifies vegetative propagation in males under conditions that would normally induce reproductive growth. The phenotype was underpinned by transcriptomic changes, showing that *MpSWI3* modulates key regulators of gametangiophore initiation (e.g. BONOBO, GLID), sperm development and motility (e.g. DUO1, PKAR), and asexual reproduction (e.g. KAI2). Together, this positions Mp*SWI3* as a chromatin-level regulator balancing vegetative and reproductive phases, highlighting an ancient epigenetic function that coordinates developmental phase transitions in land plants.

## Introduction

Land plants (Embryophyta) exhibit a haplodiplontic life cycle, in which both haploid and diploid phases are multicellular and proliferate through mitosis (Niklas and Kutschera 2010; Bowman et al. 2017; de Vries and Archibald 2018). The haploid gametophyte produces gametes mitotically, whereas the diploid sporophyte produces spores meiotically that develop into new gametophytes. This ‘alternation of generations’ contrasts with the diplontic life cycle of animals and the haplontic cycle of many streptophyte algae, where only the diploid and haploid phase, respectively is multicellular (Niklas and Kutschera 2010; Bowman et al. 2017). Nevertheless, all eukaryotic life cycles progress through an encounter and fusion of gametes that marks the onset of the diploid phase with the formation of a zygote. In most eukaryotes, including algae (Kalshoven et al. 1990; McCourt et al. 2004), this encounter depends on flagellated male gametes (sperms or spermatozoids). Following the transition of the streptophyte algal ancestor of land plants onto land, early plants still depended on flagellated spermatozoids and liquid media or a moist substrate for sperm motility and fertilization. While this is still true for the majority of land plant lineages, flagellated spermatozoids were lost twice, within conifers and angiosperms(Meyberg et al. 2020). Extant gymnosperms such as cycads produce both flagellated sperm and pollen, whereas angiosperms (flowering plants) rely entirely on pollen for fertilization (Renzaglia 2001; Niklas and Kutschera 2010). Bryophytes, which diverged from vascular plants soon after plant terrestrialization, retain flagellated spermatozoids and in contrast to spermatophytes are gametophyte (haploid) dominant(Pandey et al. 2022). As such, they provide a valuable system to study the regulation and molecular control of male reproductive development in plants(Renzaglia 2001).

Reproductive development, which mediates the transition between haploid and diploid phases through gamete fusion, is orchestrated by genetic and epigenetic regulators that likely date back to the origin of eukaryotes (Goodenough and Heitman 2014; She and Baroux 2014; Speijer et al. 2015). In plants, this regulation involves DNA-binding proteins, chromatin modifiers, and transcription factors that together form networks coordinating developmental transitions through epigenetic and protein–protein interactions (She and Baroux 2014; Borg and Berger 2015). Among these, chromatin-remodeling complexes (CRCs) form a nexus between regulatory signaling and chromatin-based transcriptional control (Clapier and Cairns 2009; Han et al. 2013). SWI3, a key subunit of the SWITCH/SUCROSE NONFERMENTING (SWI/SNF) CRC, modulates nucleosome architecture via H3 acetylation at lysine 27 (H3K27), thereby activating reproductive genes and counteracting PRC2-mediated H3K27 methylation (Sarnowski et al. 2005; Zheng and Chen 2011; Wu et al. 2012). In *Arabidopsis*, SWI3-dependent acetylation of floral identity and reproductive genes (e.g., AP3 and SEP) ensures proper flowering timing and viable seed formation (Molitor et al. 2014; Yan et al. 2019). This activity counteracts H3K27me3-mediated repression that silences key reproductive regulators such as FLOWERING LOCUS C (FLC) and maintains *AGAMOUS, APETALA3*, and *SEPALLATA* genes inactive until floral induction *(Jiang et al. 2013; Zheng et al. 2019)*.

In flowering plants, the SWI3 family has diversified into four members forming two phylogenetic groups, SWI3A/B and SWI3C/D. Likely due to an ancestral genome reduction event(Zhang et al. 2020; Donoghue et al. 2021; Harris et al. 2022; Linde et al. 2023), however, bryophytes encode only a single SWI3A/B-type gene (Genau et al. 2021). In the moss *Physcomitrium patens*, SWI3A/B mutants fail to degrade cell-wall layers during spermatogenesis, resulting in impaired sperm motility, incomplete cytoplasmic reduction, infertility, and defective male reproduction (Genau et al. 2021). While SWI3A/B function has been investigated in *Arabidopsis* and *Physcomitrium*, both species are monoecious, producing male and female reproductive organs on the same individual (Koornneef and Meinke 2010; Rensing et al. 2020). As a result, possible sex-specific roles of SWI3 in gametophyte differentiation, transcriptional regulation, and parental contribution remain unclear. *Marchantia polymorpha*, a dioecious bryophyte with sex chromosomes determining distinct male and female individuals (Bowman 2016), hence provides a system to address this issue. Furthermore, genes controlling gametangial development are only expressed during gametangiogenesis in *Physcomitrium*(Perroud et al. 2018), but they are already differentially expressed during the vegetative stage in *Marchantia*, hinting at their sex-specific regulation in the liverwort already before its reproductive organs form(Hisanaga et al. 2019).

Here, we establish that *Marchantia* SWI3 is a critical regulator in coordinating the transition from vegetative to reproductive growth in male plants and that it governs key aspects of sexual development. Remarkably, Mp*SWI3* overexpression mutants reinforce vegetative propagation instead of sexual reproduction, suggesting MpSWI3 acts as an epigenetic control point between these distinct strategies of survival and reproduction.

## Results

### SWI3 regulates reproductive development and spermiogenesis in *Marchantia*

To investigate the role of SWI3 in the haploid-to-diploid transition and sexual development, we attempted to generate distinct male and female lines with mutations in *Marchantia SWI3*, using CRISPR/Cas9(Sugano et al. 2018). sgRNAs targeting the *MpSWI3* coding region never yielded any viable transformants, but sgRNA targeting the 5’ upstream region provided us with male and female mutants carrying insertions, deletions, or substitutions in their promoter (Fig. S1a). Quantitative real-time PCR showed all lines to overexpress *MpSWI3* in comparison to the WT (Fig. S1b), which is why we refer to them as *swi3*OE-♂ and *swi3*OE-**♀** hereafter. The vegetative growth was unaffected in both the *swi3*OE lines (Fig. 1a). After two weeks under far red light – which induces the transition to reproductive phase – *swi3OE*-♂, however, produced approximately half as many antheridiophores (male gametangiophores) as the wild type (WT). In contrast, archegoniophore (female gametangiophores) production in the mutants was comparable to the WT (Fig. 1a-b). These results established *Mp*SWI3 as a gametangiophore development regulator in males.

**Fig. 1:**
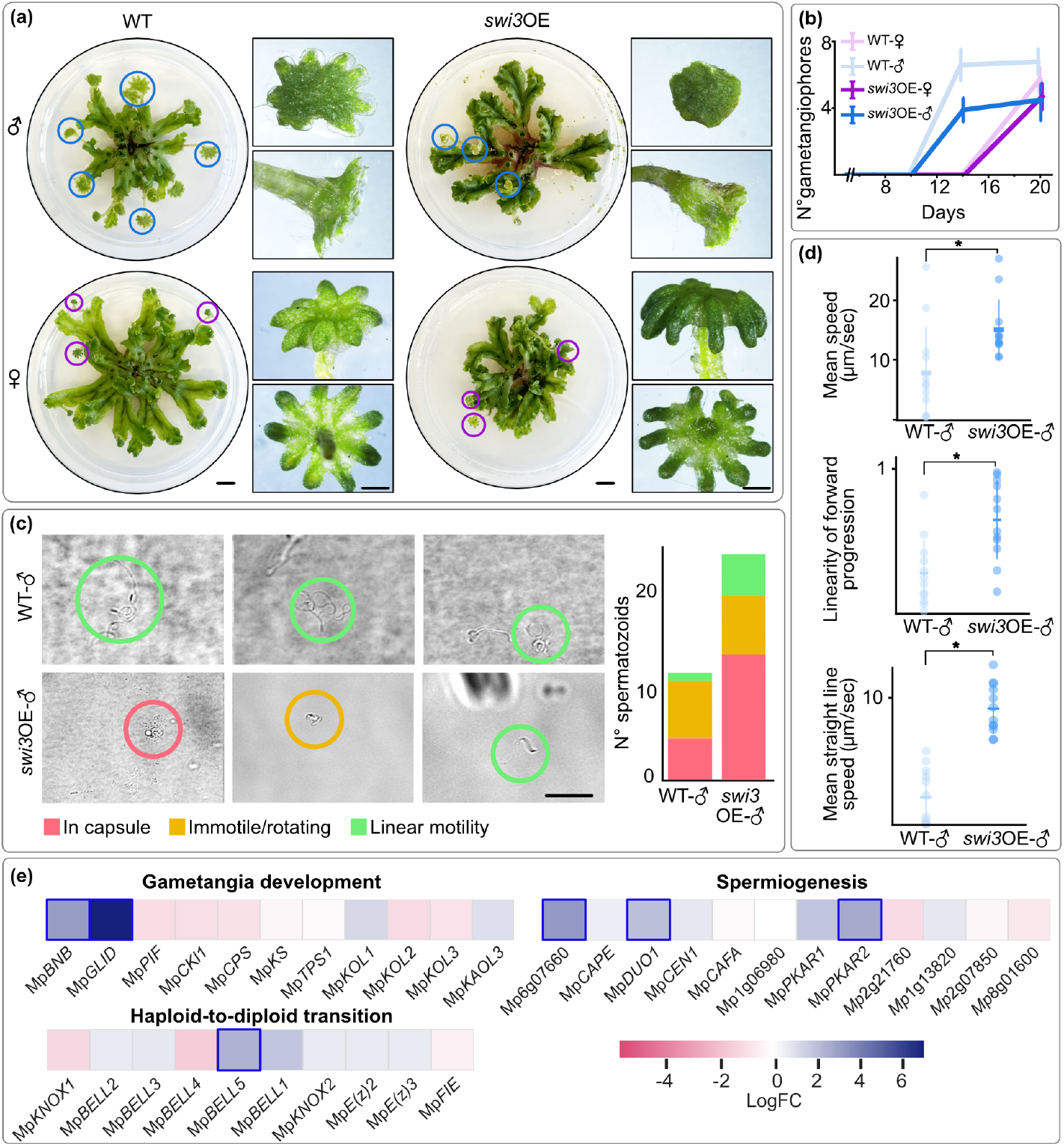
*Mp*SWI3 mutants show defective antheridiophore formation and altered spermiogenesis. **(a)** Representative 30 days-old male and female plants from wild-type and *swi3*OE lines. Full view of the thalli with their gametangiophores highlighted by circles, which are shown enlarged in the squares. **(b)** Gametangiophore maturation over time (n=15). **(c)** Representative images of sperms (left) and sperm motility patterns (right) from WT-♂ and *swi3*OE*-*♂. Scale bar: 20μm. **(d)** Mean speed, linearity of forward progression, and mean straight line speed of sperms from WT and *swi3*OE-♂ lines (n=10). Statistically significant differences (P value < 0.05) are indicated by asterisks. **(e)** LogFC values for a key subset of genes differentially expressed in *swi3*OE-♂ as compared to the WT-♂ under far-red light conditions; genes with significant differences in their expression are marked by a blue outline.

Given the role of *Mp*SWI3 in antheridiophore development, we next examined whether *swi3*OE-♂ antheridiophores produced motile sperms. WT spermatozoids matured to stage-5 as expected, whereas those of *swi3*OE-♂ frequently arrested at the earlier stages (Fig. 1c). In WT, 38.5% of sperm remained capsulated, whereas in *swi3*OE-♂, this proportion increased to 55.6% (Fig. 1c). *Swi3*OE-♂ sperms also exhibited a significantly higher straight-line speed, greater linearity of forward progression, and overall increased swimming velocity (Fig. 1d). These results indicate that *Mp*SWI3 influences both early spermiogenesis and motility dynamics of mature sperms.

To uncover the molecular basis of SWI3’s role in regulating reproductive development and spermiogenesis in male *Marchantia* plants, we performed comparative RNA-seq analysis of plants grown under far-red light. *Swi3*OE-♂ showed an upregulation of 584 genes and a downregulation of 409 (|log_2_FC| > 2) (Fig. S2a, Table S1). Functional annotation of differentially expressed genes revealed enrichment for categories related to plant defence, peroxidase activity, and cell-wall remodelling (Fig. S2b). We next examined genes specifically linked to sexual and asexual reproduction. Among those involved in gametangiophore and gametangia development, Mp*BONOBO* (Mp*BNB*) and Mp*GLID* were strongly upregulated (log_2_FC > 3), whereas Mp*PIF*, Mp*CKI1*, Mp*CPS*, and Mp*KOL2/3* were downregulated (Fig. 1e). Genes associated with the haploid-to-diploid transition showed a mixed pattern: Mp*BELL1* and Mp*BELL5* were upregulated, while Mp*KNOX1* and Mp*BELL4* were downregulated. Transcripts linked to sperm differentiation and motility, including Mp*DUO1* (a master regulator of spermiogenesis in land plants (Higo et al. 2018)), Mp*PKAR* (a regulator of sperm motility(Yamamoto et al. 2024)) and a flagellar radial spoke protein homolog (Mp6g07660), were upregulated (Fig. 1e, Table S2). These patterns are consistent with the morphology and motility changes observed in *swi3*OE-♂ sperms in *Marchantia* (Fig. 1c–d) as well as *Physcomitrium*, where SWI3a/b loss of function led to impairment in late maturation(Genau et al. 2021).

### SWI3 overexpression reinforces vegetative growth *in lieu* of reproductive development

Because *Mp*SWI3 influenced both gametangiophore development and sperm maturation under far red light, we examined whether it also affected the vegetative to reproductive transition under far red light. WT plants showed the expected morphological responses (e.g. thallus flattening, gametangiophore formation, and empty gemma cups; gemma are vegetative propagules). However, *swi3*OE-♂ produced significantly more gemma cups (16.2 ± 1,5) than WT-♂ (10.5 ± 1,2), whereas *swi3*OE -♀ and WT-♀ showed no difference (Fig. 3a–b). Interestingly, while WT and *swi3*OE-♀ suppressed gemmae production, *swi3*OE-♂ produced 13.5-fold more gemmae at day 10 (Fig. 2). These results demonstrate that *Mp*SWI3 overexpression promotes a shift to asexual reproduction through excessive gemma production under far-red light, instead of sexual reproduction via gametangiophore development. This shift was facilitated by expression changes in a number of genes involved in vegetative growth (Fig. 2d). This included the substantial overexpression of *Mp*KAIB, a key gemmae-formation factor contributing to the excessive gemma formation in *swi3*OE-♂(Komatsu et al. 2023) (Fig. 2). Moreover, several bHLH transcription factors (*Mp*SETA, *Mp*BHLH29, *Mp*BHLH30, *Mp*BHLH31, *Mp*BHLH21, and *Mp*PIF) were downregulated, whereas late embryogenesis-related genes such as Mp*LEA-like44* and Mp*LEA-like45* were strongly upregulated (Fig. S2c). Together, these transcriptional changes suggest that *Mp*SWI3 may regulate genes related to sexual differentiation as well as asexual propagation, positioning it as a chromatin-level switch in the liverwort, balancing reproductive and vegetative growth, rather than unidirectional control of reproductive onset.

**Fig. 2:**
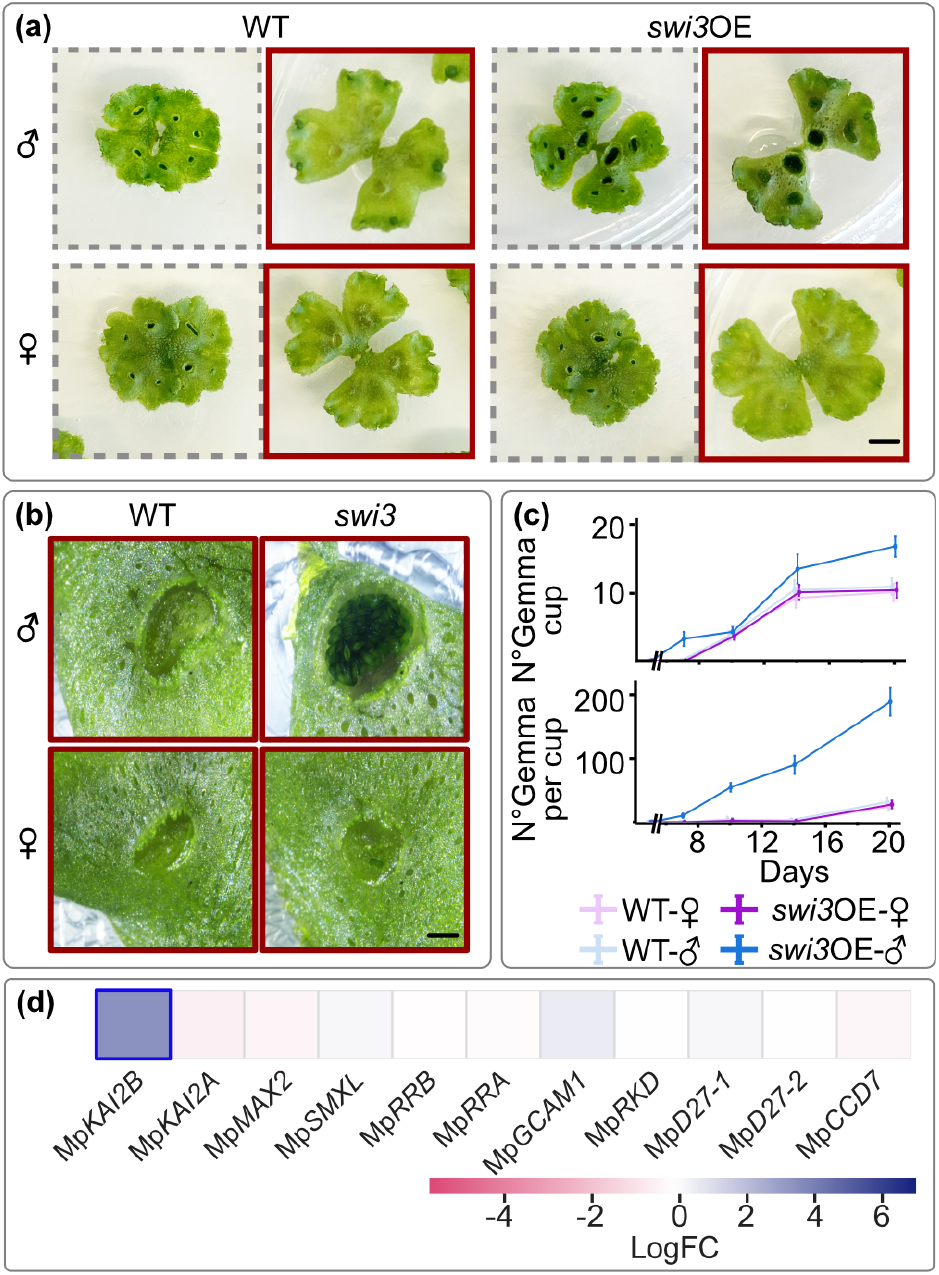
*swi3*OE-♂ overproduce gemmae under far-red light. **(a)** Representative plants of 14d-old wild-type and *swi3*OE lines. Plants outlined with a grey dashed border were grown under continuous white light, whereas those outlined in red were supplemented with far-red light. Scale bar: 0.4cm. **(b)** Closer view of representative gemmae cups of plants grown with supplemented far-red light. Scale bar: 0.2cm. **(c)** Gemma cup formation over time and number of gemmae per cup counted; n=15 for thalli and cups analysed. **(d)** LogFC values for key genes involved in gemma formation in *swi3*OE-♂ (as compared to the WT-♂) under far-red light conditions. Genes with significant differences in expression are marked by a blue outline.

## Discussion

Sexual reproduction is a defining trait of eukaryotic life, ensuring genetic recombination through the fusion of haploid gametes to form a diploid zygote(Goodenough and Heitman 2014; Speijer et al. 2015; Garg and Martin 2016; Raval et al. 2022; Jeffries et al. 2025). Across eukaryotic lineages, this fundamental process is elaborated by mechanisms that regulate the differentiation, compatibility, and union of gametes – from mating-type systems in unicellular algae and fungi to complex sexual organs in multicellular plants and animals(Heitman 2015; Yadav et al. 2023; Rizos et al. 2024; Becker et al. 2025). This diversification accompanied the transition from aquatic to terrestrial habitats, where new reproductive structures and regulatory cues evolved to ensure gamete encounter and fertilization in plants(Niklas and Kutschera 2010; She and Baroux 2014; Mori et al. 2015; Becker et al. 2025). These reproductive structures range from specialized gametangia to the evolution of pollen and seeds, which together enable fertilization independent of a wet habitat. Soon after the algal transition to land, and prior to the extensive diversification of reproductive structures seen in vascular plants, the ancestor of all bryophytes emerged after genome reformatting(Zhang et al. 2020; Donoghue et al. 2021; Harris et al. 2022; Linde et al. 2023). Consequently, extant bryophytes allow to explore the emergence of developmental and regulatory complexity underlying sexual differentiation, and the transition between vegetative and reproductive phases from a different perspective.

Much of our understanding of the regulatory networks governing the transition between vegetative and reproductive phases, however, comes from studies in angiosperms. Here, numerous transcription factors, signalling cascades, and chromatin-remodelling complexes have been characterized(Blazquez & Weigel 2000; Andrés and Coupland 2012; He 2012). A key role for SWI3 is evident in *Arabidopsis*, where it regulates multiple reproduction-related developmental traits, including flowering time and reproductive organ formation(Sarnowski et al. 2005). Loss of *At*SWI3A or *At*SWI3B results in embryo abortion and male sterility, respectively, while mutations in *At*SWI3C or *At*SWI3D alter sexual-organ development(Sarnowski et al. 2005). Our findings extend this role to the bryophyte *Marchantia*, establishing *Mp*SWI3 as a key regulator of the vegetative–reproductive balance in a sister lineage to seed plants.

Sexual reproduction in *Marchantia* is induced through the coordinated action of light-signalling pathways that integrate far-red irradiance and photoperiodic cues(Inoue et al. 2019). Key genes include Mp*BNB* and Mp*GLID*, which drive the formation of male gamete precursor cells(Ren et al. 2024). In *swi3*OE-♂, both these genes are significantly upregulated, yet the plants developed fewer gametangiophores. It suggests that *Mp*SWI3 overexpression may cause premature or ectopic activation of these two genes in vegetative tissues, producing – as previously reported(Ren et al. 2024) – a transcriptional signature of reproductive induction without proper organ differentiation. Such an uncoupling between germline specification and organogenesis in *swi3*OE*-♂* likely reflects impaired FR-phytochrome-PIF signalling and reduced gibberellin (GA) biosynthesis, consistent with the observed downregulation of Mp*PIF*, Mp*CPS*, and Mp*KOL*. PIF- and GA-dependent pathways act upstream of the Mp*BNB–*Mp*GLID* module to promote organ initiation and maturation; their downregulation would therefore weaken gametangiophore formation, even when germline markers remain elevated(Sun et al. 2023). Similarly, GA-deficient mutants produce fewer and morphologically abnormal gametangiophores(Sun et al. 2023). Together, these observations suggest that *SWI3* overexpression disrupts the coordination between germline activation and organ differentiation.

Beyond its role in male reproductive development, our results uncover a striking reversal of developmental fate, as excess SWI3 reinforces vegetative propagation through excessive gemmae production in *swi*3OE-♂ under conditions that normally induce reproductive growth. A key regulator of gemmae production and vegetative growth is *Mp*KAI2, which drives gemma cup formation and gemma initiation(Komatsu et al. 2023). Strong overexpression of *Mp*KAI2B in *swi3*OE-♂ lines underlies this reversal towards vegetative growth despite reproduction-inducing cues. This finding highlights a broader role for SWI3 in balancing vegetative and reproductive programs as an epigenetic modulator of phase transitions in the bryophyte. Such a shift toward vegetative propagation, particularly in lines where reproductive growth is compromised, may reflect a bet-hedging strategy in which vegetative reproduction ensures survival when sexual reproduction is impaired. This regulatory flexibility could allow plants in nature to abort or delay reproductive development, when conditions are suboptimal for fertilization, maintaining persistence through vegetative growth. Such environmentally driven developmental plasticity might have been crucial for the survival of early land plants adapting to novel terrestrial stresses and continues to contribute to the resilience of extant bryophytes that thrive across diverse habitats.

Altogether, our findings position SWI3 as a key chromatin regulator orchestrating the balance between vegetative propagation and sexual reproduction in *Marchantia* males. This is the first report of sex-specific roles of SWI3, and during the vegetative stage (before gametaniophore formation), in contrast to other species (Genau et al. 2021) (Sarnowski et al. 2005). By modulating the accessibility of key regulatory loci, *Mp*SWI3 likely integrates environmental and developmental cues to determine whether cells commit to vegetative or reproductive fates. These results underscore that epigenetic mechanisms were essential for synchronizing gene expression with developmental contex during the origin of embryophytes. On the molecular level, SWI3 via modulation of H3 K27 acetylation probably acts as a counterpart of PRC2, a writer of H3K27me3 repressive marks(Sarnowski et al. 2005; Zheng and Chen 2011; Wu et al. 2012). Factors such as far red light apparently influence this balance, and an unbalance such as by the overexpression in *swi3*OE-♂ is able to tip the system into different states. Understanding how SWI3-mediated chromatin remodeling governs these ancestral phase transitions reveals the molecular basis of reproductive plasticity in bryophytes and provides first evolutionary insight into how chromatin dynamics and diversification of factors such as SWI3 facilitated the shift from reversible vegetative–reproductive balance in early land plants to fixed floral commitment in angiosperms. The mechanisms by which chromatin regulation integrates environmental and developmental signals to determine reproductive fate remain poorly understood outside of angiosperms. Bryophytes, particulary dioecious species, provide a powerful model to uncover how these ancestral processes evolved into the complex reproductive strategies of terrestrial plants.

## Materials and methods

### Plant growth conditions

*Marchantia polymorpha* (BoGa ecotype; Botanical Garden, Osnabrück University, Germany) was cultivated on half-strength Gamborg’s B5 medium containing 1% glucose and 1% agar under continuous full light spectrum (450-700nm) 70 μmol m^-2^ s^-1^ at 22°C. To induce the reproductive stage, the light conditions were supplemented with far-red light (700-750nm) 40 μmol m^-2^ s^-1^ at 22°C in ECO^2^ boxes (oval, 80 mm) with a green filter #40 (Duchefa Biochemie).

### Generation of mutant plants

CRISPR/Cas9-based genome editing of MpSW3a/b (Mp8g15610) was performed as previously described (Sugano et al. 2018). Three sgRNAs (sgRNA1,2,3) were designed, sgRNA1 targeted the promoter region, sgRNA2 and sgRNA3 targeted the coding sequence of *Mp*SWI3 (Fig. S1). Cloning of sgRNA was performed using pMpGE_En03 as the entry- and pMpGE010 as a destination vector using LR Clonase II enzyme mix (Thermo Fisher Scientific). Destination vectors were introduced into electrocompetent *Agrobacterium tumefaciens* (GV301 without pSOUP) by electroporation (Bio-Rad GenePulser Xcell, 1.44 kV). *Agrobacterium*-mediated sporeling transformation was performed as described previously(Ishizaki et al. 2008). After three days of spore co-cultivation with *Agrobacterium* transfectants, sporelings were washed three times with liquid ½ Gamborg’s B5 medium to remove Agrobacteria and plated on agar plates with half-strength Gamborg’s B5 medium with 1% Glucose supplemented with 10 μg/mL Hygromycin and 100 μg/mL Cefotaxime for selection.

### Macroscopic phenotyping

To assay gametangiophore numbers, 15 plants per line were grown in ECO^2^ boxes that allow the upwards growth of gametangiophores. Each gametangiophore and gemma cup was counted as soon as it could be seen by sight from the top of the thallus. To quantify gemmae per cup, all gemmae from gemmae cups closer to the thallus base were harvested by a toothpick, gemmae were spread on a plastid sheet, imaged and counted using ImageJ(1.54f).

### Sperm phenotyping

Sperms were harvested by applying water on top of the antheridia from 30 days old plants cultivated under far red light. After 5 min of incubation, the water was collected and sperm morphology monitored with a Nikon eclipse Ti imaging platform. For quantitative motility analysis, sperms were video-recorded at the resolution of 1280 x 1024 pixels and at the rate of 33 frames per second (fps) by a Nikon Digital Sight DS-U3. Movies were converted into a sequence of individual frames and sperm movement tracked by the ImageJ plugin TrackMate v7.11.1. Two movies were analyzed to obtain sperm swimming motility parameters for each line.

### RNA-seq

Total RNA was extracted from WT and *swi3*OE-♂ plants incubated under far red light for 14 days, using the RNeasy Plant Mini Kit for RNA Extraction (Qiagen). Three biological replicates were prepared for each genotype and mRNA isolated via their poly(A) tail. RNA integrity and concentration were assessed using a NanoDrop spectrophotometer. Libraries were prepared by Eurofins Genomics (Ebersberg, Germany). Sequencing was performed on an Illumina NovaSeq 6000 platform, generating paired-end reads (2 × 150 bp), with an average of ∼30 million read pairs per sample. Sequencing data was quality checked and analyzed as described previously(Frangedakis et al. 2024). Briefly, high quality reads were retained using FASTQC and TrimGalore, aligned to *Marchantia polymorpha* genome (v.7.1) using Kallisto(Bray et al. 2016; Bowman et al. 2017) and DEG were obtained through DESeq2(Love et al. 2014).

## Data availability

Supplementary figures and tables are available with this submission. Transcriptome data are available in the NCBI Sequence Read Archive: PRJNA1353859.

## Author contributions

MLQ: supervision; experimental design, investigation, data curation, formal analysis, validation and visualization, writing original draft, review and editing. PKR: formal analysis, visualization, writing original draft, review and editing. SR: conceptualization; funding and resource acquisition. SBG: conceptualization; project administration; supervision; experimental design; funding and resource acquisition; data visualization; writing original draft, review and editing.

## Acknowledgments

We acknowledge support from the high-performance computing cluster (HILBERT) and thank Daniel Wasim Djamriani, Lilly Möbus, Franklin Cooper, and Margarete Stracke for their help. We also thank Sabine Zachgo and Nora Gutsche from the University of Osnabrück (Germany) for providing us with spores, and Zhanghai Li for providing us with pMpGE010_sgRNA1. PKR is grateful to Bill Martin for providing financial support.

