## Supplementary figures for "SWI3 regulates male sex determination in *Marchantia polymorpha*"

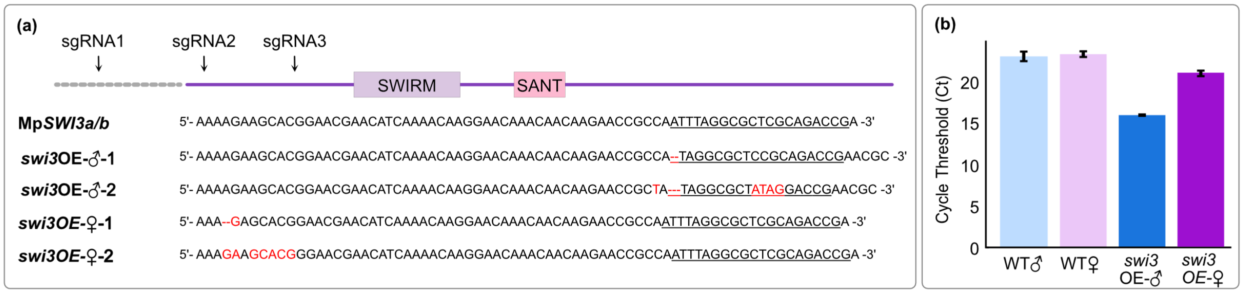


**Fig. S1: Confirmation of *Mp*SWI3 mutants. (a)** Sequencing results of the targeted Mp*SWI3* promoter region from the wild type (WT) and Mp*SWI3* mutants. **(b)** Quantitative real-time PCR of two week old thalli.

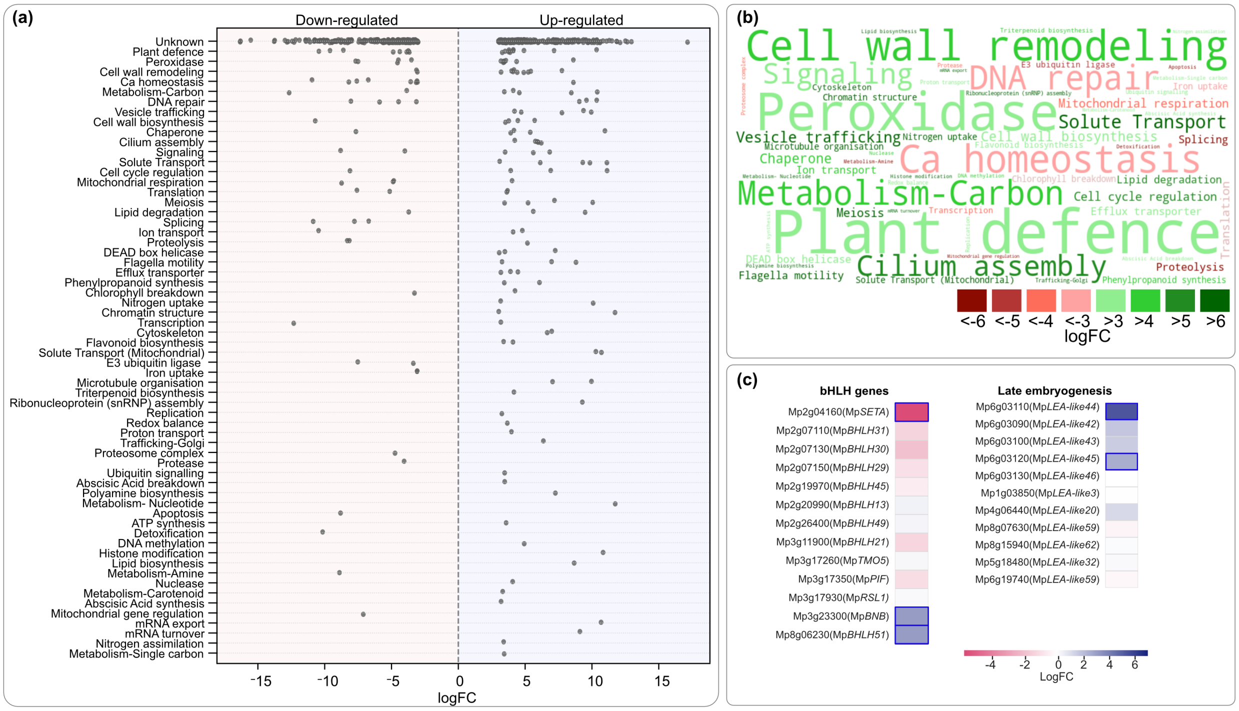


**Fig. S2: Global gene expression changes in *swi3*OE-♂ under FR light. (a)** 30% of the differentially expressed genes (DEGs; $Log_2FC>3$, P value<10e-6) could be functionally annotated using the Kyoto Encyclopedia of Genes and Genomes (KEGG) database. Each row of the scatter plot indicates a functional category, and each data point is a DEG under that category. The genes on the right (purple background and positive logFC values) are upregulated and the genes on the left are downregulated in *swi3*OE-♂. **(b)** Functional categories of DEG are shown in a word cloud, where the word size indicates the number of DEGs under that category (i.e. a bigger word size indicates a higher number of DEGs under that category). The color indicates the median LogFC change of the DEGs under that category as per the color key on the bottom right. **(c)** logFC values for a select subset of DEG; genes with significant differences in regulation are marked by cells with a blue outline.
